# Investigating the effect of POPC-POPG Lipid Bilayer Composition on PAP248-286 Binding using CG Molecular Dynamics Simulations

**DOI:** 10.1101/2023.07.20.549823

**Authors:** Nikhil Agrawal, Emilio Parisini

## Abstract

PAP248-286 is a fusogenic peptide derived from prostatic acid phosphatase, commonly found in human semen, and is known to mediate HIV fusion with cell membranes. In this study, we performed 120 independent coarse-grained molecular dynamics simulations to investigate the spontaneous binding of PAP248-286 monomers, considering both charged and neutral histidine (His) residues, to membrane bilayers composed of different lipid compositions: 100% POPC, 70% POPC-30% POPG, and 50% POPC-50% POPG. Our simulations revealed that PAP248-286 displayed spontaneous binding to the membrane, with increased binding observed in the presence of anionic lipid POPG. Specifically, in systems containing 30% and 50% POPG lipids, monomer residues, particularly in the systems containing charged histidine (His) residues, exhibited prolonged binding with the membrane. Furthermore, our simulations indicated that PAP248-286 adopted a parallel orientation with the membrane, exposing its positively charged residues to the lipid bilayer. Interestingly, systems containing charged His residues showed higher lipid occupancy around the peptide. These findings are consistent with previous experimental data, suggesting that PAP248-286 binding is enhanced in membranes with charged His residues, resembling the conditions found in the acidic vaginal pH environment. The results of our study provide further insights into the molecular mechanisms underlying the membrane binding of PAP248-286, contributing to our understanding of its potential role in HIV fusion and infection.

## 1. Introduction

Human immunodeficiency virus (HIV) infection poses a major global health challenge^1^. According to the WHO 2023 report, more than 84 million people have been infected by HIV and more than 40 million people have lost their lives to the virus^2^. Sexual transmission continues to be one of the primary modes of HIV transmission, with human semen serving as a vector for viral transmission, and women, in particular, at higher risk of contracting HIV through sexual contact^3-4^. One factor that has drawn the interest of the HIV research community is the role of a specific 39 amino acid-long peptide, PAP248-286, in the infection process^5-6^. PAP248-286 is derived from prostatic acid phosphatase (PAP), a protein present in high concentrations in human seminal fluid^7^. Previous studies have suggested that PAP248- 286 peptides play an important role in enhancing HIV infection by several folds^5, 8^. PAP248- 286 peptides aggregate to form amyloid fibrils termed semen-derived enhancers of viral infection (SEVI)^5, 9-10^. Unlike typical amyloid peptides, which mainly display biological activity when aggregated into fibrils, PAP248-286 has been reported to show biological activity even in its monomeric form^5, 11-15^.

The interaction between amyloid monomers and cell membranes can alter the integrity and functionality of the membranes, producing cytotoxic effects^16-17^. Conversely, membranes can also expedite the aggregation and fibril formation process of these peptides, thereby further enhancing membrane disruption^18-20^. The highly cationic nature of the SEVI fibrils facilitates their binding to the anionic membranes of the HIV virions. This binding interaction promotes the bridging between the viral and host cell membranes, leading to an increased propensity for viral attachment and subsequent infection^10,21-22^. In addition to its role in enhancing HIV infection, SEVI fibrils have demonstrated the ability to facilitate infection by cytomegalovirus, herpes simplex virus types 1 and 2, and Ebola virus. Furthermore, SEVI fibrils have been implicated in sperm quality control mechanisms and have been observed to agglutinate bacteria^23-27^. While the biological activity of PAP248-286 has primarily been attributed to the SEVI amyloid, it is noteworthy that even in its monomeric form (i.e., without assembly into SEVI fibrils), PAP248-286 exhibits the capability to enhance HIV infection in the presence of seminal fluid^5^.

In a previous experimental study Brender *et. al*.^28^ showed that PAP248-286 monomers bind with the membrane in α-helical conformation and suggested that the peptide binds to the membrane in a parallel orientation. This binding configuration plays a crucial role in promoting bridging interactions between membranes. The study suggested that the parallel binding of the peptide in its α-helical conformation could serve two important functions: i) dehydrating the lipid headgroups, thereby facilitating fusion, and ii) enabling positively charged residues, such as lysine, to interact with lipid headgroups and effectively reduce the long-range coulombic repulsion between the bilayers. In their study they also found that PAP248-286 induced greater lipid aggregation in liposomes containing 7:3 POPC/POPG lipids at an acidic pH resembling the vaginal environment (pH ∼4.2), compared to neutral pH. This effect was attributed to the two histidine residues in PAP248-286, which are likely to be charged at the acidic pH but not at neutral pH. In another experimental study Nguyen *et. al*.^29^ utilized Sum Frequency Generation vibrational spectroscopy (SFG) to explore the interaction between PAP248-286 peptide and lipid bilayers composed of 7:3 POPC/POPG and 7:3 DPPC/DPPG. Their investigations unveiled that PAP248-286 peptide exhibited a more pronounced and intimate interaction with the 7:3 POPC/POPG lipid bilayer when compared to its gel-phase counterpart. Furthermore, their study revealed that the structure of PAP248-286, when bound to lipids, adopts an α-helical conformation, resembling its secondary structure in a 50% solution of 2,2,2-trifluoroethanol.

In the present study, we conducted a total of 120 independent coarse-grained molecular dynamics (MD) simulations of PAP248-286 with bilayers containing different compositions of lipids. The aim of this study is to examine the interaction of PAP248-286 monomer in two scenarios: one with charged His residues representing the vaginal environment, and the other without charged His residues representing the semen environment. These monomers were examined in bilayers with different compositions, including 100% POPC, 70% POPC-30% POPG, and 50% POPC-50% POPG. A total of 120 microseconds (μs) long simulations were performed with each individual run lasting for 1 μs. Through these simulations, we were able to investigate how different concentrations of POPG in the bilayer influence the spontaneous binding of PAP248-286 monomer to the membrane. Additionally, we identified the key amino acids involved in the binding process and determined the occupancy and lipid distribution around PAP248-286 during the simulations.

## 2. Methods

### 2.1. Structures and force field of PAP248-286 fibrils and POPC-POPG membranes

For the CG molecular dynamics simulations, we utilized the NMR structure of a PAP248-286 monomer (PDB id: 2L77 and BMRB Entry: 17346) as the starting model^30^. The residue numbering in the PDB id: 2L77 structure ranges from 1 to 39. At pH 4.2, His3 and His23 of PAP248-286 are protonated, resulting in an overall peptide charge of +8, while at pH 7.2, the charge is +6. To account for the acidic vaginal environment and the physiological environment, we used the H++ server^31^ to add hydrogen atoms to PAP248-286 at pH 4.2 and pH 7.2, respectively. The atomistic structures of both the PAP2486-286 with protonated His residues and PAP248-286 with deprotonated His residues were converted into a CG model (Fig. 1) using the CHARMM-GUI Martini maker^32-33^. Membrane bilayers of different compositions (100% POPC, 70% POPC-30% POPG and 50% POPC-50% POPG) were prepared using the CHARMM-GUI membrane bilayer builder option. Table 1 shows the initial number of lipid molecules in each type of membrane. Prior to starting the simulations on the fibril-membrane complexes, membranes were equilibrated for 100 ns each and the average area per lipid was calculated for the last 20 ns for each simulation. For 100% POPC average area was 65.49 ± 0.92 Å^2^, for 70% POPC-30% POPG average area was 64.70 ± 1.00 Å^2^, and for 50% POPC-50% membrane average area was 64.42 ± 0.89 Å^2^.

**Table 1.** Number of lipids in each leaflets.

| Membrane bilayer | Number of POPC molecules | Number of POPG molecules |
| --- | --- | --- |
| 100% POPC | 200 (100 in each leaflet) | 0 |
| 70% POPC-30% POPG | 140 ( 70 in each leaflet) | 60 (30 in each leaflet) |
| 70% POPC-50% POPG | 100 (50 in each leaflet) | 100 (50 in each leaflet) |

**Fig. 1.**
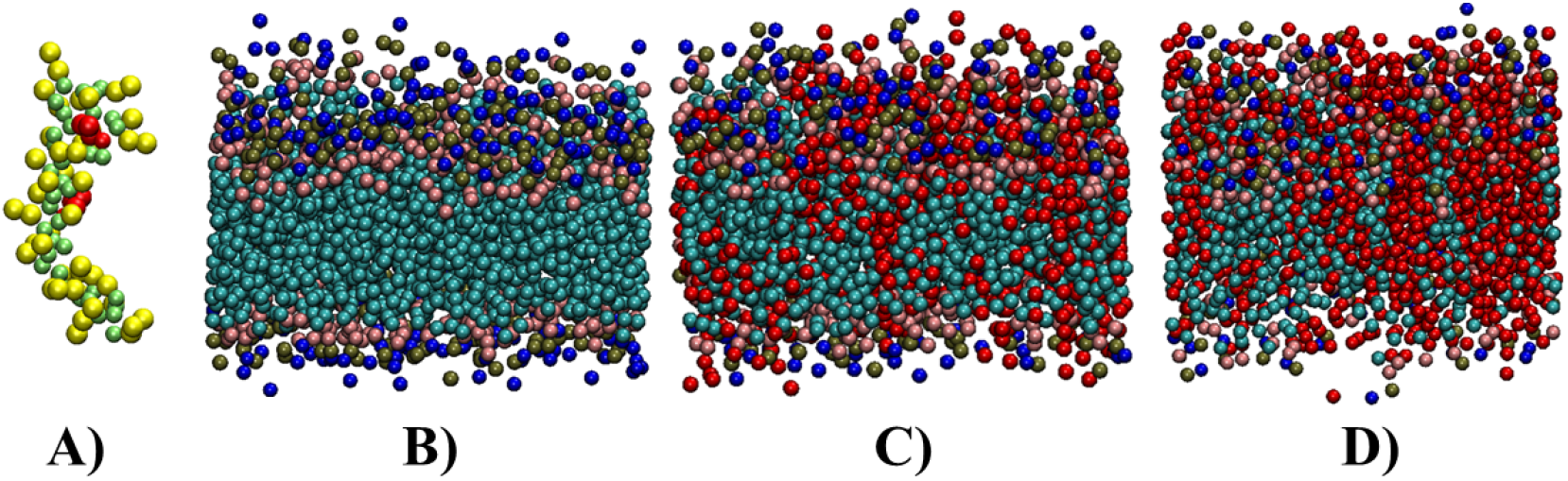
A) CG model of the PAP248-286 peptide (PDB id: 2L77), with the two histidine residues highlighted in red, B) 100% POPC membrane bilayer. D) 70% POPC-30% POPG membrane bilayer, C) 50% POPC-50% POPG membrane bilayer. CG model of peptide is shown using VdW representations, while backbone beads of peptide are shown in light green colour, side chain beads are shown in yellow colour. The head group beads NC3 and PO4 of the POPC molecules are illustrated in blue and tan and the tails are depicted in cyan, respectively, while POPG lipids are shown in red colour.

We employed the Martini 2.2 protein force field^34^ to simulate the PAP248-286 peptide, and for the membrane, water, and ions, we utilized the Martini 2.0 force field^35^.

### 2.2. Simulation protocol

In our study, we examined a total of six PAP248-286-membrane systems. The first three systems consisted, each, of one PAP248-286 monomer with charged His residues. System one featured a 100% POPC bilayer, 9201 water molecules, and 8 CL^-^ ions for system neutralization. System two involved a 70% POPC-30% POPG bilayer, 8913 water molecules, and 52 NA^+^ ions for system neutralization. System three comprised a 50% POPC-50% POPG bilayer, 8812 water molecules, and 92 NA^+^ ions for system neutralization. Additionally, we studied three systems where the PAP248-286 monomer had no charged His residues. System four consisted of a 100% POPC bilayer, 9172 water molecules, and 6 CL^-^ ions for system neutralization. System five involved a 70% POPC-30% POPG bilayer, 8910 water molecules, and 54 NA^+^ ions for system neutralization. Lastly, system six comprised a 50% POPC-50% POPG bilayer, 8789 water molecules, and 94 NA^+^ ions for system neutralization. Initially, all six systems underwent energy minimization using the steepest descent algorithm^36^.All systems underwent equilibration in five cycles, with decreasing restraints applied in each cycle. The force constants used were 1000 kJ mol^-1^ nm^-2^ in the first cycle, decreasing to 50 kJ mol^-1^ nm^-2^ in the last cycle for the PAP248-286 peptide, and 200 kJ mol^-1^ nm^-2^ in the first cycle, decreasing to 10 kJ mol^-1^ nm^-2^ in the last cycle for the lipid head groups. The equilibration process lasted a total of 4750 ps. During the production simulations, no restraints were applied to the PAP248-286 peptide and membranes. The Parrinello-Rahman algorithm^37^ with semi-isotropic pressure coupling was used for pressure coupling, and the velocity-rescale algorithm^38^ was used for temperature coupling. Pressure coupling was performed with a bath time of 12.0 ps, and temperature coupling with a bath time of 1.0 ps. All simulations were conducted at a temperature of 310.15 K and a pressure of 1 atm. Newton’s equations of motion were integrated using a time step of 20 femtoseconds (fs). A cut-off distance of 1.1 nm was applied for Van der Waals (VdW) and electrostatic interactions, with the reaction field method used for the treatment of electrostatic interactions. A total of 120 independent simulations were carried out, with each simulation lasting 1 μs, and the initial random velocities and all simulations were performed using the GROMACS 2021 simulation package^39^.

### 2.3. Analysis details

To determine the minimum distance between the PAP248-286 peptide and the membrane, we utilized the GROMACS minidist program^39^. For interaction analysis, we considered an interaction between the PAP248-286 peptide residue and the membrane when the minimum distance between any bead of the membrane lipids (POPC or POG) was ≤LJ5LJÅ of any beads of the PAP248-286 peptide. These calculations were performed for the whole simulation time. Additionally, we calculated the number of lipid molecules within 5 Å of PAP248-286 using GROMACS select program and lipid occupancy was determined by counting the frames where there were more than 0 lipid molecules within the 5 Å proximity of the membrane, divided by the total number of frames. The area per lipid was calculated using FATSLiM^40^.

## Results

### 3.1. Binding of the PAP248-286 with the membranes

PAP248-286 is a fusogenic peptide known to disrupt the integrity of cell membranes, potentially enhancing fusion mediated by the HIV gp41 protein^11, 28^. To investigate the spontaneous binding of PAP248-286 monomers (with charged and neutral His residues) to the membrane, we analysed the time evolution of the minimum distance between PAP248-286 and the membrane in various systems (Fig. 2). A trajectory was considered a binding event if the PAP248-286 peptide interacted with or remained bound to the membrane for at least 200 nanoseconds (ns). Out of 40 independent trajectories of PAP248-286 with a 100% POPC bilayer, we observed binding in 13 trajectories (8 trajectories with charged His and 5 trajectories with neutral His, as shown in Fig. 2A and 2D). In the case of PAP248-286 with bilayers containing 70% and 30% POPG, we observed binding in 33 out of 40 trajectories (17 trajectories with charged His and 16 trajectories with neutral His, Fig. 2B and 2E). Similarly, in the case of bilayers containing 50% POPC and 50% POPG, binding was observed in 39 out of 40 trajectories (20 trajectories with charged His and 19 trajectories with neutral His, Fig. 2C and 2F). These findings suggest that an increased presence of POPG, which leads to a more anionic membrane, significantly enhances the binding of PAP248-286. Furthermore, the data indicates that PAP248-286 with charged His residues exhibits a higher number of binding events in 100% POPC systems, whereas there is no substantial difference in systems containing POPG lipids.

**Fig. 2.**
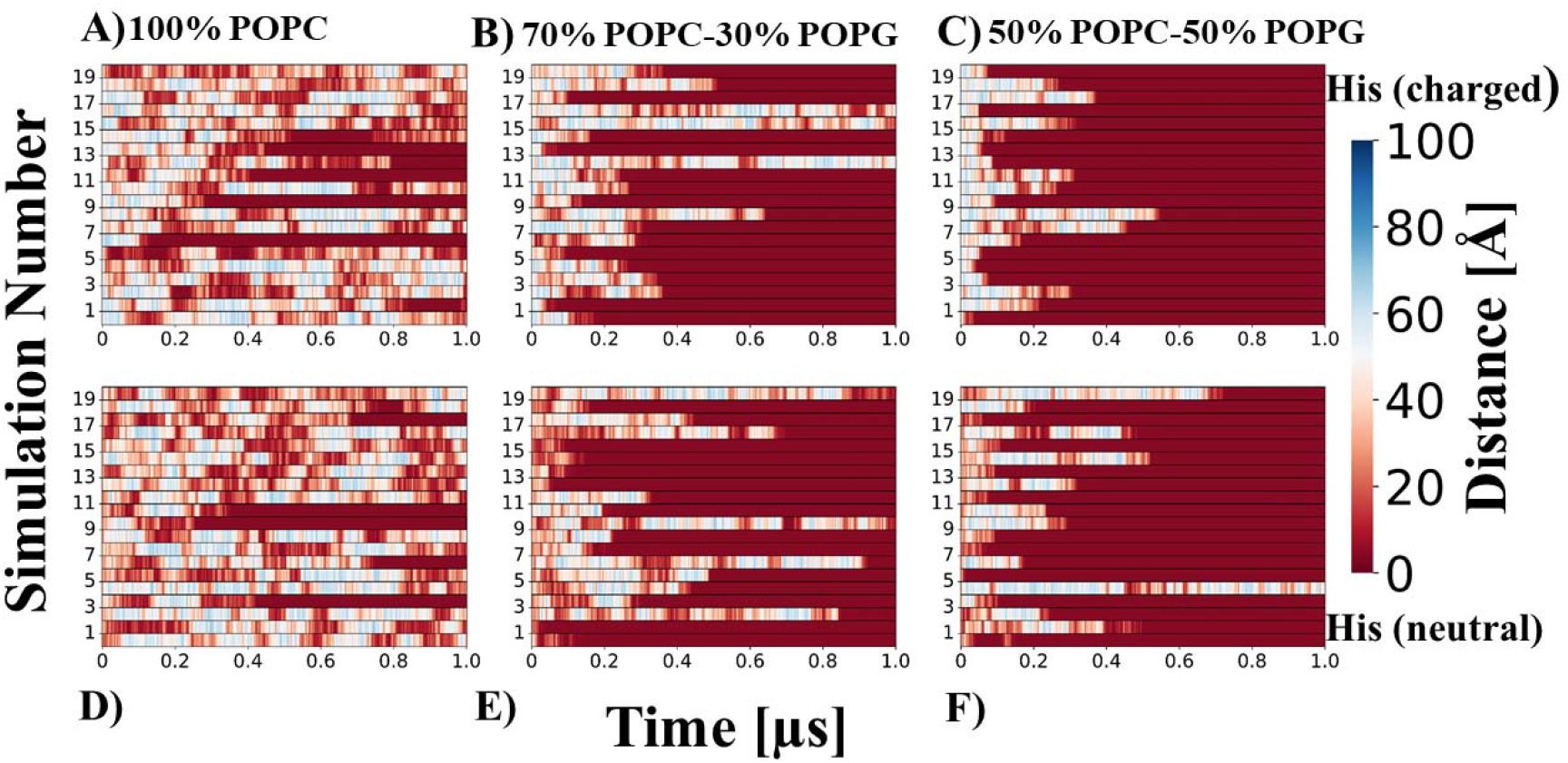
A-C) Time evolution of the minimum distance between PAP248-286 (His residues charged) and 100% POPC, 70% POPC-30% POPG and 50% POPC-50% POPG membranes, respectively. D-F) Time evolution of the minimum distance between PAP248-286 (His residues neutral) and 100% POPC, 70% POPC-30% POPG and 50% POPC-50% POPG membranes, respectively.

The orientation of PAP248-286 is believed to play a significant role in its interaction with the membrane and its ability to facilitate HIV infection^5, 28^. To gain further insight into this mechanism, we captured snapshots of PAP248-286 peptide with the membrane at the initial and final time points from a representative trajectory (Fig. 3), specifically from systems with 70% POPC-30% POPG and 50% POPC-50% POPG compositions. Our observations revealed that PAP248-286 peptide binds parallel to the membrane surface, adopting a “carpet pose” orientation. It does not insert into the membrane but rather remains positioned atop the membrane. This arrangement allows the peptide to expose all of its residues to the membrane, potentially facilitating interactions with the membrane components. By adopting a parallel orientation without membrane insertion, PAP248-286 maximizes its contact with the membrane, enhancing the potential for interactions and influencing the membrane’s integrity.

**Fig. 3.**
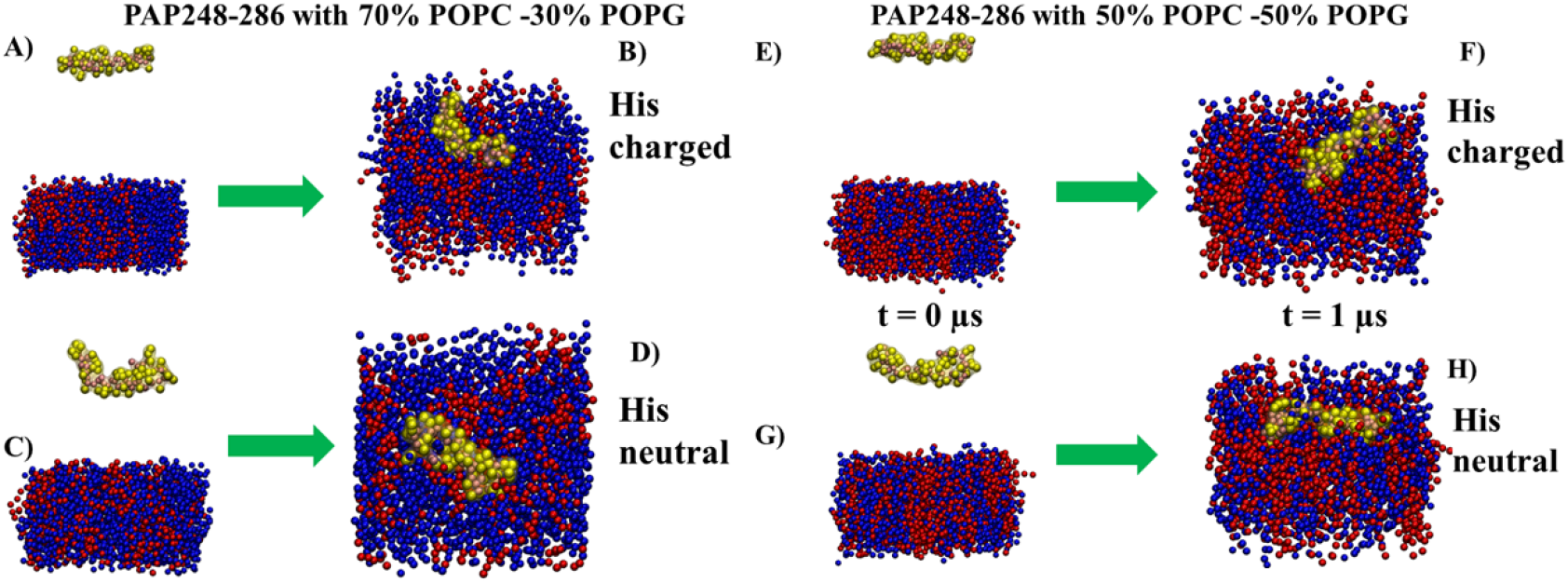
Structures of the PAP248-286 (HIS residues charged and neutral) with membrane containing 70% POPC and 30% POPG, and with membrane containing 50% POPC and 50% POPG at two different time points from four representative trajectories. A-B) Representative image for PAP248-286 (His residues charged) with 70% POPC-30% POPG at 0 and 1 μs. C-D) Representative image for PAP248-286 (His residues neutral) with 70% POPC-30% POPG at 0 and 1 μs. E-F) Representative image for PAP248-286 (His residues charged) with 50% POPC-50% POPG at 0 and 1 μs. C-D) Representative image for PAP248-286 (His residues neutral) with 50% POPC-50% POPG at 0 and 1 μs. PAP248-286 peptide, and membranes are shown in VdW representation. The backbone beads of PAP248-286 peptide are depicted in pink, and the side chains beads are shown in yellow. The POPC membrane has been depicted in blue colour and POPC and POPG in red colour.

### 3.2. identification of residues involved in the binding of PAP248-286 to the membranes

To determine the residues that bind to the membrane, we analyzed the simulation data by calculating the percentage of time each residue remained within 5 Å of the membrane for at least 50% of the simulation duration (Fig. 4). We then identified key binding residues as those that bound to the membrane in at least 80% of the simulations across multiple independent trajectories. In the simulations of PAP248-286 with a membrane composed of 100% POPC, most residues showed binding in less than 50% of the simulations, except in 4 out of 40 trajectories. In these specific trajectories, residues Tyr33, Lys35, Leu36, Ile37, Met38, and Tyr39 interacted with the membrane for at least 50% of the simulation time. However, no single residue was consistently identified as a key binding residue across all simulations. For PAP248-286 peptide simulations with a membrane consisting of 70% POPC and 30% POPG, we observed binding of residues for at least 50% of the simulation time in 26 out of 40 trajectories (15 trajectories with charged His residues and 11 trajectories with neutral His residues). In the simulations with charged His residues, residues His3, Lys35, and Leu36 consistently bound to the membrane for at least 80% of the simulation time across multiple trajectories. In the simulations with neutral His residues, we did not observe any residues that consistently bound for at least 80% of the simulation time in multiple trajectories. In addition to these residues, two binding patches were identified: residues 1-4 (Gly1, Ile2, His3, Lys4) and residues 35-39 (Lys35, Leu36, Ile37, Met38, Tyr39), which interacted with the membrane for at least 50% of the simulation time in most trajectories, with His23 also showing particularly strong binding. In simulations of PAP248-286 with a membrane composed of 50% POPC and 50% POPG, residues bound for at least 50% of the simulation time in 35 out of 40 trajectories (19 trajectories with charged His residues and 16 trajectories with neutral His residues). Two major binding regions were identified: residues 1-4 and 34-39, which consistently showed strong binding. Additionally, residues Ser9, Arg10, Leu16, Val17, His23, and Met24 also exhibited significant interaction with the membrane. In the simulations with charged His residues, residues Ile2, His3, His23, Lys35, and Leu36 showed interactions with the membrane for at least 80% of the simulation time in multiple trajectories. In the simulations with neutral His residues, Tyr33 and Lys35 interacted with the membrane for at least 80% of the simulation time. Overall, these findings suggest that a greater number of residues interacted with the membrane for longer durations in systems containing charged His residues in the presence of 30% POPG and 50% POPG in the PAP248-286 peptide.

**Fig. 4.**
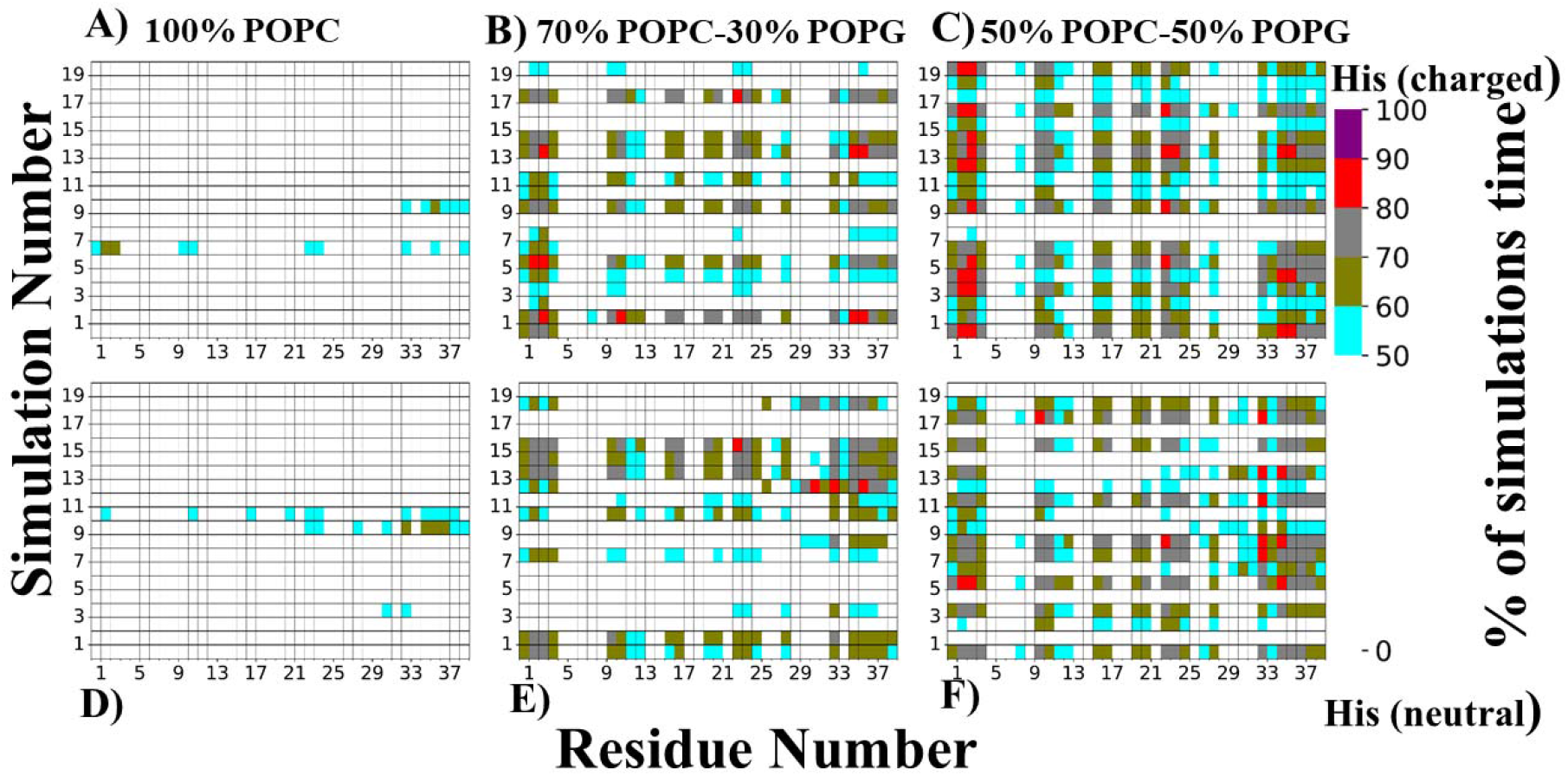
Percentage of time in which residues of the PAP248-286 (His charged) remained at a distance ≤□5□Å from the membrane A) 100% POPC, B) 70% POPC-30 % POPG, C) 50% POPC-50% POPG. Percentage of time in which residues of the PAP248-286 (His neutral) remained at a distance ≤□5□Å from the membrane A) 100% POPC, B) 70% POPC-30 % POPG, C) 50% POPC-50% POPG.

### 3.3. Lipid occupancy and number of lipids within 5Å of PAP248-286 peptide

We caculated lipid occupany (see method section of more details) in all simulations trajectoies, and to futher see distribution of number of lipids in all the systems we plotted boxplots. The data (Fig.5 A) revealed a significant increase in lipid occupancy around the PAP248-286 peptide as the concentration of POPG lipids in the membrane increased. Furthermore, when comparing the systems containing PAP248-286 with charged His residues and neutral His residues, we observed higher lipid occupancy in the charged His residues systems. This indicates that the presence of charges on the His residues facilitated increased interaction between PAP248-286 and lipid molecules. To examine the distribution of the number of lipids in all the systems, we constructed boxplots (Fig. 5). In the 100% POPC bilayer systems, there was not much difference in the number of lipids between the His charged and His neutral systems. However, in the systems containing 70% POPC and 30% POPG, we observed a range of lipid numbers from 0 to approximately 8 in the His charged system, with a median value of around 5 and the highest number of lipids reaching approximately 18. In the His neutral system, the number of lipids ranged from 0 to around 6, with a median of approximately 4 and the highest number of lipids around 17. The most notable disparity was observed in the systems containing 50% POPC and 50% POPG. In these systems, the His charged system had lipid numbers ranging from approximately 4 to 10, with the highest number of lipids reaching up to 20. Conversely, in the His neutral system, the number of lipids ranged from 0 to approximately 7, with the highest number of lipids around 18. Overall, this data suggests that there were more lipids surrounding His charged residues systems containing 30% and 50% POPG.

**Fig. 5.**
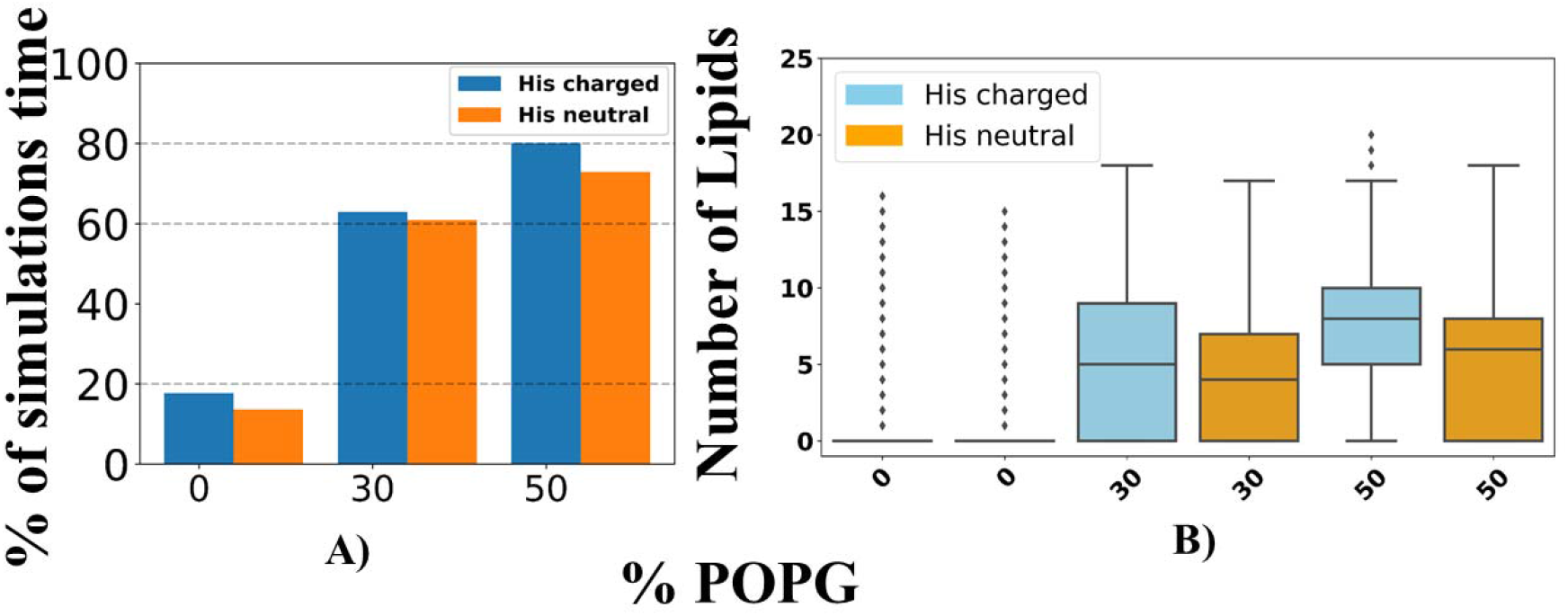
A) Lipid occupancy in all three systems containing no POPG, 30% POPG and 50 % POPG, B) Boxplots of number lipids of all three systems containing no POPG, 30% POPG and 50 % POPG.

## 4. Discussion & conclusions

In our study, we conducted coarse-grained molecular dynamics (MD) simulations to investigate the interaction between the PAP248-286 peptide and membrane bilayers. The bilayer was composed of different concentrations of POPG lipids: no POPG, 30% POPG, or 50% POPG. Our findings revealed that the binding of the PAP248-286 peptide to the membrane increased with an increase in the concentration of POPG lipids. Additionally, we observed that the PAP248-286 peptide with charged His residues exhibited a higher number of binding trajectories in all three systems. Moreover, in the systems with 30% POPG and 50% POPG, the PAP248-286 peptide with charged His residues showed a greater number of residues remaining within 5Å of the membrane. These systems also demonstrated increased lipid occupancy and a higher number of lipids surrounding the PAP248-286 peptide with charged His residues compare to systems containing neutral His. We also observed that PAP248-286 peptide prefers to bind with membranes in the parallel position or in “carpet pose”, which allows the peptide to expose most of its residues to the membranes, leaving the other side available to bind the other membrane, which might be helpful in fusion activity. Although binding was non-specific, we saw that some key residues such as His3, His23, Tyr33, Lys34, Lys35 showed highest proximity to the membranes, especially in the systems containing the charged His residues.

The findings from our study align with previous research^28^ that investigated the binding behaviour of PAP248-286 peptide to the membrane. It was previously suggested that PAP248-286 binds to the membrane in a parallel orientation, allowing its positively charged residues to be exposed to the membrane. Our data supports this notion and further confirms that the binding of PAP248-286 to the membrane is influenced by the charged state of the histidine residues. In particular, our results indicate that PAP248-286 exhibits higher binding affinity to the membrane when the histidine residues are charged, such as at acidic pH. This is consistent with previous studies that have highlighted the amphiphilic and electrostatic nature of histidine, which allows it to effectively reside at membrane interfaces. Histidine’s ability to engage in both hydrophobic and electrostatic interactions contributes to its role in membrane binding^41-42^. Similarly, lysine residues have been recognized as important mediators of membrane perturbation due to their involvement in electrostatic interactions with negatively charged phospholipid head groups and hydrophobic interactions with the lipid hydrocarbon chains. The characteristics of lysine residues make them effective agents in disturbing membranes^43-44^. Overall, our data supports the idea that the presence of POPG lipids in the membrane enhances the binding of PAP248-286 peptide, particularly when the peptide contains charged histidine residues. Furthermore, our observations indicate that systems with higher concentrations of POPG lipids promote closer proximity of peptide residues to the membrane and a greater number of lipid molecules in the vicinity of the peptide. This underscores the significance of lipid composition in modulating the interaction between the peptide and the membrane. In conclusion, our findings contribute to a deeper understanding of the peptide-membrane interaction and highlight the importance of charged residues, such as histidine and lysine, in mediating the binding of membranes.

## Acknowledgments

N.A. acknowledges the ERDF postdoc grant (No. 1.1.1.2/VIAA/4/20/757) for project funding. E.P. thanks the ERDF project BioDrug (No. 1.1.1.5/19/A/004) and the Latvian Council of Science (grant No. lzp-2020/2–0013) for financial support. We would like to thank Latvian Institute of Organic Synthesis for computational resources. N.A. would also like to thank the Centre for High Performance Computing (CHPC) in Cape Town (South Africa) for supercomputing resources.

